# A stochastic compartmental model to simulate intra- and inter-species influenza transmission in an indoor swine farm

**DOI:** 10.1101/2022.11.18.517057

**Authors:** Eric Kontowicz, Max Moreno-Madriñan, Darryl Ragland, Wendy Beauvais

## Abstract

Common in swine production worldwide, influenza causes significant reductions in feed efficiency and potential transmission to the workforce. Swine vaccines are not universally used in swine production, partly due to their limited efficacy because of continuously evolving influenza viruses. We evaluated the effects of vaccination, quarantine of infected pigs, and changes to workforce routine (ensuring workers moved from younger pig batches to older pig batches). A Susceptible-Exposed-Infected-Recovered model was used to simulate stochastic influenza transmission during a single production cycle on an indoor hog growing unit containing 4000 pigs and two workers. The absence of control practices resulted in 3,958 pigs [1 – 3972] being infected and a 0.61 probability of workforce infection. Quarantine of infected pigs the same day they became infectious was the single effective control practice, reducing the number of infected pigs to 3 [1 - 3961] and the probability of workforce infection to 0.27. The second-best control practice was mass pig vaccination (80% effective vaccine), which reduced the number of infected pigs to 23.5 [1 - 635] and the probability of workforce infection to 0.07. The third-best control practice was changing the worker routine by starting with younger to older pig batches, which reduced the number of infected pigs to 997 [1 - 1984] and the probability of workforce infection (0.22). All other control practices, when considered alone, showed little improvement in reducing total infected pigs and the probability of workforce infection. Combining all control strategies reduced the total number of infected pigs to 1 or 2 with a minimal probability of workforce infection (0.03 – 0.01). These findings suggest that non-pharmaceutical interventions can reduce the impact of influenza on swine production and reduce the risk of interspecies transmission when efficacious vaccines are unavailable. Therefore, these results can help prevent the emergence of influenza strains with pandemic potential.

## Introduction

Influenza A viruses (IAV), members of the Orthomyxoviridae family, are common respiratory viruses that infect humans and animals, and have a worldwide distribution (1). Domestic pigs *(Sus domesticus*) play a role in the maintenance of influenza viruses globally (especially subtypes H1N1, H3N2, and H1N2 within the United States). Influenza viruses associated with swine can spill over to humans or other species. In most cases, the virus is not well adapted to transmission via the new host, however host adaptation can evolve via point mutations and genetic reassortment. The risk that this process leads to a pandemic is difficult to quantify but is non-negligible (2-13). In 2009 a novel subtype of H1N1 IAV spilled over from swine to humans leading to a pandemic which was estimated to infect approximately 60.8 million Americans (14), and one to three billion people worldwide (15-45% of the world’s population) (15). It is likely that swine as well as avian hosts also played a role in the evolution of the highly virulent influenza virus responsible for the 1918 pandemic, which killed 50-100 million people in a time when the world population was 1.7 billion (16).

In both pigs and humans, influenza virus is mainly transmitted via inhalation of fomites and aerosols that are generated through breathing, coughing or sneezing (17). Additionally, indirect contact with fomites on contaminated equipment or worker clothing have also been implicated as routes of transmission for IAV in swine populations (17). In some cases, IAV can replicate in the intestinal tract of pigs, shed in feces, and in turn transmit to a susceptible host (18, 19). On typical US swine farms, morbidity can be extremely high (up to 100% of the population) while mortality is typically low (< 1%) (20). However, in naïve populations, mortalities can exceed 10% of the infected population (21). Infected pigs are estimated to lose 5 to 12 pounds in body weight over a three-to-four-week period (22-24), leading to prolonged finishing times which contribute to the economic burden that influenza places on swine producers.

In addition to economic impacts, swine-associated influenza may also be transmitted to the swine workforce. One study found swine workers’ odds of elevated antibodies to swine H1N1 virus (A/swine/WI/238/97) to be 54.9 (95% confidence interval [95% CI]13.0-232.6) times those of non-swine industry worker controls (25). In a separate seroprevalence study evaluating farmers, meat processing workers (from pork production facilities), veterinarians with swine exposure and controls with no known contact with swine, it was found that farmers had the highest odds of swine influenza exposure (Odds Ratio [OR], 35.3; 95% CI, 7.7 – 161.8) (26). Another study using whole genome sequencing found that IAV samples from five workers had identical clade classification to swine-origin IAV samples from pigs on the same farm (27).

Evidence also suggests that IAV can be transmitted from workers to pigs. One study in the Czech Republic found the presence of antibodies in pigs to human-origin influenza (6). In the US, transmission of 2009 H1N1 influenza from workers back into the pig population has frequently been recorded (28). Some influenza viruses (e.g., A/California/VRDL101/2009/H1N1) transmitted from humans to pigs have not been shown to be transmitted within pig populations while other influenza viruses (e.g. A/swine/Illinois/A01395201/2013/H1N1) transmitted by humans can become well established in pigs as determined via phylogenic relationships (28). Reverse zoonotic transmission (human-to-animal transmission) of influenza virus leading to sustained swine-to-swine transmission have been reported relatively infrequently; 20 such events were reported between 1965 and 2013 (6). However, it is highly likely that transmission events have gone undetected.

### Control measures

Currently, vaccination is one of the most practical and effective means to control and prevent influenza in pigs and humans alike. However vaccine efficacy is frequently suboptimal and limited by the rapid evolution of influenza viruses (29, 30). Whole, inactivated viruses with adjuvant are typically administered via intramuscular injection to either growing pigs or sows (31, 32). Maternally-derived antibodies have been shown to protect offspring for approximately 6 weeks post-weaning (33). Of the commercial inactivated vaccines used in the US, it is estimated that around 50% are autogenous (30) *i*.*e*. subtypes isolated from swine farms are used directly to make a vaccine for use in the same population (34). In the US, veterinarians may request authorization to use these autogenous vaccines from the Food and Drug Administration. In addition to inactivated vaccines, viral vector vaccines, live attenuated vaccines, DNA vaccines and subunit IAV vaccines are commercially available but used less frequently (30, 34). Reduced viral shedding due to vaccination has been documented in both humans (35) and pigs (36-38).

In addition to vaccination, grower farms may practice all-in all-out practices (with thorough cleaning and disinfection between batches), biosecurity practices such as limiting personnel and showering in and out, purchasing piglets from known sources, surveillance of incoming animals, isolation and quarantine (17, 39). Ventilation systems have been shown to reduce the entrance of and transmission of pathogens across different rooms on farms (40-43). Lastly, improved personal protective equipment (PPE) or adherence to PPE mandates can help reduce influenza transmission between hogs and workers. Despite these available control measures, IAV remains prevalent in US grower herds (44). One study of an animal operation in Minnesota found that specialized footwear was the most commonly used form of PPE by workers and they were unlikely to use masks or gloves (45). This study also found that handwashing before and after animal handling was not widely encouraged (45).

Etbaigha et al. (2016)(46) adapted a deterministic model first published by Reynolds et al. (2014)(47) to show that piglets play a key role in the maintenance of IAV on farrow-to-finish farms. Reynolds further showed that implementing either homologous or heterologous mass vaccination was not sufficient to eliminate IAV on a wean-to-finish farm (47). Using a stochastic approach, both Pitzer et al. (48) and White et al. (49) indicated that piglets are a key reservoir for IAV and vaccination alone will not eliminate the presence of IAV on breeding and finishing farms. Further, White et al. (49) showed that a combination of vaccination and biosecurity practices can reduce the likelihood of maintaining an endemic reservoir in piglets on a breeding farm.

The goal of this study was to quantify the effectiveness of IAV control measures targeting both hogs and the workforce within a single, typical U.S. indoor hog-growing unit. The current model differs from previous research by: (1) incorporating stochastic transmission between pigs and the workforce, (2) evaluating multiple combinations of prevention, biosecurity and workplace policies on IAV dynamics within a US indoor growing unit, and (3) comparing the effectiveness of these measures in terms of cases averted in both pigs and the workforce.

## Methods

### Overview

We simulated stochastic transmission of a single influenza virus subtype amongst swine and the workforce during one production cycle. We extended the compartmental modelling approach used by Pitzer et al (48) and White et al (49) by adapting the swine population model to a typical US hog growing production unit and including interspecies transmission between the workforce and swine. We first tested a baseline model that assumes current typical industry practices and no vaccination. We then explored transmission dynamics under different intervention strategies to evaluate the change in the number of pigs infected, the probability of at least one workforce infection, and the time until interspecies transmission (to workers or pigs). Intervention strategies included mass vaccination (40% effectiveness or 80% effectiveness), quarantine of sick pigs, changes to worker routine and improved PPE including masks.

### Population Dynamics

Two populations were simulated: the growing pig population and the human workforce on the farm (**Figure 1**). The farm layout, number of pigs per room, and total number of workers employed on the farm closely resemble a typical commercial hog grower unit in Indiana and other swine intensive areas in the Midwest. These details were determined via informal expert consultation.

**Figure 1.**
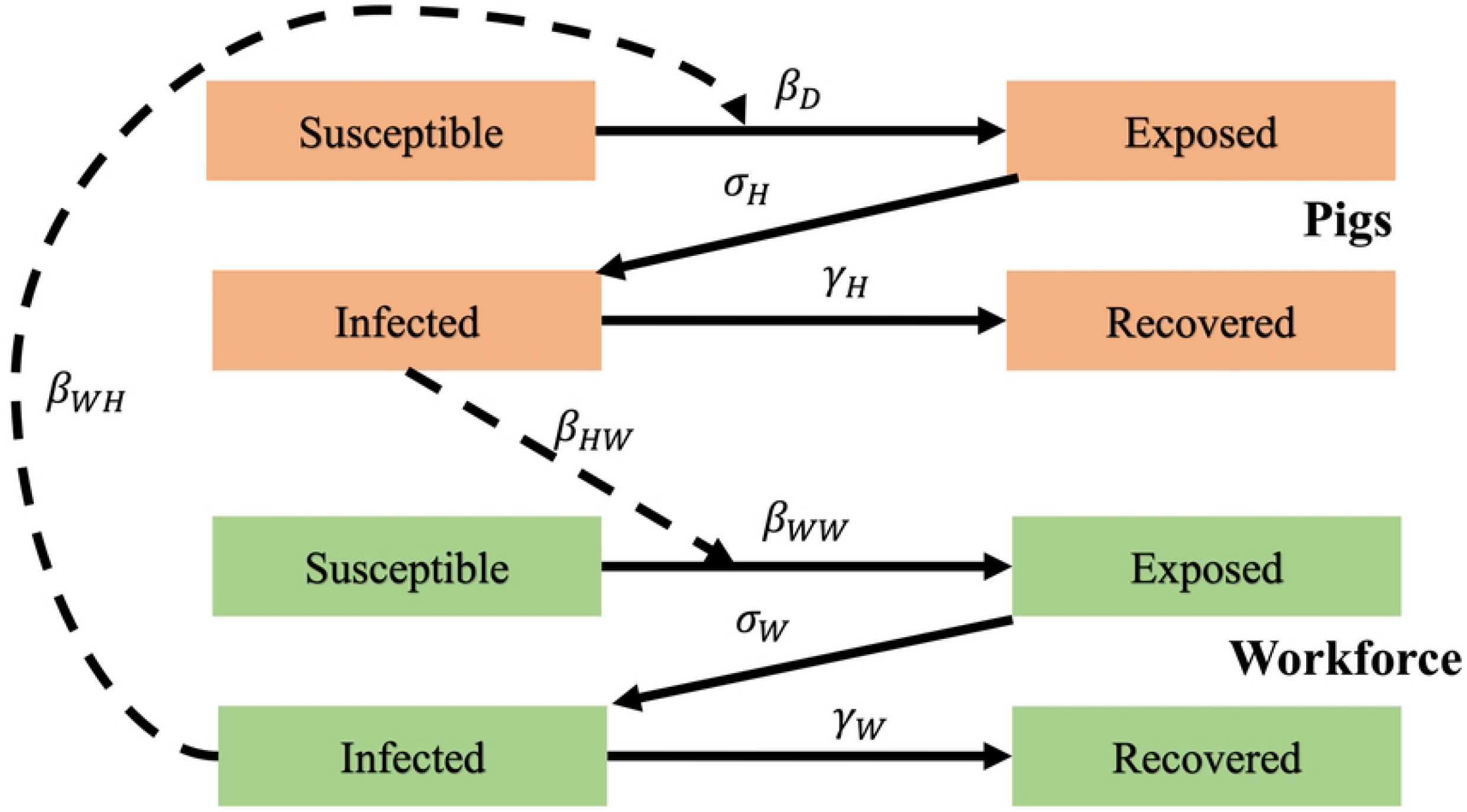
Compartmental model to predict influenza transmission dynamics on a typical US hog grower unit. We assume hogs entered at 3 weeks of age and remained for 23 weeks until they reached market weight. Each room houses 1,000 hogs and there is no direct contact between hogs in separate rooms, but indirect transmission can occur via aerosol and fomite transmission (e.g. contaminated equipment, worker clothes or boots). The workforce consists of two workers that visit each room daily and have an equal likelihood of contacting any hog.

Pigs enter the grower facility at three weeks of age after they have been weaned at a breeding and farrowing facility. When entering the grower unit, each batch of weaned pigs are placed into a single room and remain in this room for 23 weeks, when it is assumed that they have reached market weight. We assumed an all-in / all-out system (50) where each room was filled two weeks apart to maximum capacity (1,000 hogs per room; 4,000 hogs in total). All pigs in each room leave the farm on the same day. For example, room 1 is filled to 1,000 pigs on day 1, room 2 filled on day 14, etc. There is no direct contact between pigs in separate rooms. It was assumed that only two workers were needed for husbandry tasks, such as feeding and performing daily checks (50, 51). It was assumed that each worker had an equal probability of interacting with each hog.

### Transmission Dynamics

The transmission of IAV on this growing farm is simulated using a Susceptible-Exposed-Infected-Recovered (SEIR) model. *S*_*i*_(*t*), *E*_*i*_(*t*), *I*_*i*_(*t*), and *R*_*i*_(*t*) are the number of susceptible, exposed, infected and recovered pigs respectively; *t* is time in days, where t ≥ 0, *i* represents the room where the growing hog is housed, where *i* is equal to 1, 2, 3 or 4, *j* represents the adjunct rooms where other hos are housed, where *j* is equal to 1, 2, or 3. *S*_*W*_(*t*), *E*_*W*_(*t*), *I*_*W*_(*t*), and *R*_*W*_(*t*) are the corresponding compartments representing the workforce. Pigs become infected via within-room or between-room transmission. Between-room transmission occurs via air flow and movement of fomites such as contaminated workers’ clothing, boots or equipment.

Here we describe the baseline model that assumes typical control measures are used. The SEIR models for the hogs and workforce are defined by the following set of differential equations:

#### Hogs

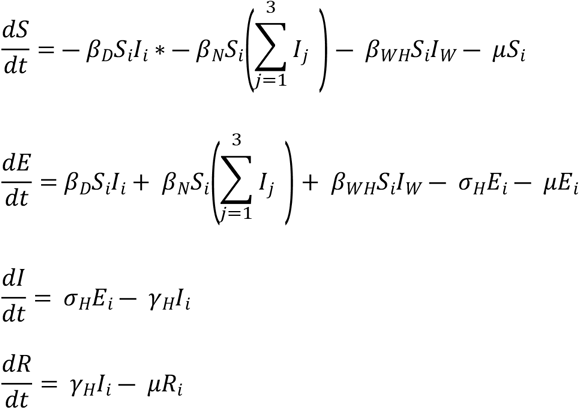

#### Workforce

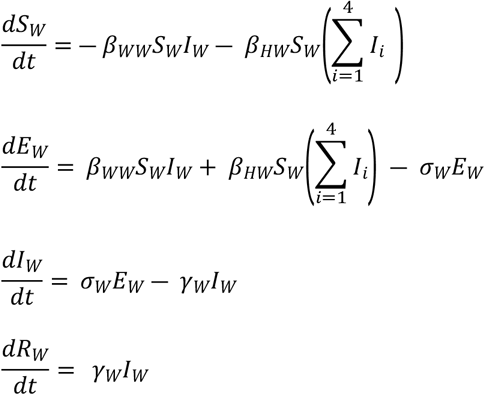

The parameters are described in **Table 1**. We assume no between-room transmission from swine to the workforce or between-room transmission between workforce members. We did not include a loss of immunity term in our model because we focused on single subtypes of influenza and single production cycles with a length of 23 weeks. Evidence suggests that natural infection with IAV can induce lasting protection against the same subtype (52). Stochastic transitions between compartments were simulated using the Gillespie Algorithm (53), using the most likely parameter values presented in **Table 1** as the mean transition rates. Distributions of parameter values were used for later sensitivity analysis (54).

**Table 1.** Parameters for IAV SEIR model with description and values.

| Parameter Description | Parameter Symbol | Most Likely Value<br>(Parameters for<br>Uncertainty<br>Distribution) | Uncertainty<br>Distribution<br>(for<br>sensitivity<br>analysis) | Source |
| --- | --- | --- | --- | --- |
| Within-room<br>Transmission Rate | $\beta_D$ | 0.002 (0.001 – 0.1) /day | Triangle | (36) |
| Between-room<br>Transmission Rate | $\beta_N$ | $\beta_d/178$<br>(0, $\beta_d$) | Uniform | (36, 55) |
| Workforce to Pig<br>Transmission Rate | $\beta_{WH}$ | 0.00007 / day (0.000001,<br>0.0001) | Uniform | (56) |
| Pig to Workforce<br>Transmission Rate | $\beta_{HW}$ | 0.00003 / day (0.000001,<br>0.0001) | Uniform | (57) |
| Quarantined Pig to Workforce Transmission Rate | $\beta_{QHW}$ | 0.0 / day | Fixed | NA |
| Workforce to Quarantined Pig Transmission Rate | $\beta_{WQH}$ | 0.0 / day | Fixed | NA |
| Workforce to Workforce Transmission Rate | $\beta_{WW}$ | 0.32 / day<br>(0.0000001, 0.64) | Triangle | (58) |
| Latent Period for Pigs | $\sigma_H$ | 2 days (sd = 1) | Normal | (36) |
| Infectious Period for Pigs | $\gamma_H$ | 5 days (sd = 1) | Normal | (36) |
| Latent Period of Humans | $\sigma_W$ | 2 days (sd = 1) | Normal | (58) |
| Infectious Period for Humans | $\gamma_W$ | 3 days (sd = 1) | Normal | (58) |
| Natural Death Rate for Pigs | $\mu$ | 0.00028 / day | Fixed | (46) |
| Quarantine Rate | $q$ | 1, 0.5, or 0.0 per infected hog / day | Fixed | NA |
| Pig Vaccine Efficacy | $Vacc\ Eff$ | 80%, 40%, or 0% | Fixed | (59-61) |

Assuming density-dependent transmission, we estimated that : *β*_*D*_ = *R*_0_/(*N* * *D*) (62), where *N* is the number of pigs in a single room (1,000 hogs), *D* is the duration of infectiousness (5 days) and *R*_*0*_ = 10, based on experimental data using a triple reassortant H1N1 subtype (A/swine/IA/00239/04) (36). We found a paucity of empirical data quantifying interspecies transmission rates within indoor hog growing units. We estimated that *β*_*HW*_ = *R*_0_/(*N* * *D*) (62), where *N* = 4002, *D* = 5 and *R*_*0*_ = 0.6, based on an outbreak of influenza A (H3N2) among attendees of an agricultural fair (57). We assumed that during this outbreak, a single pig was responsible for transmitting to the first three confirmed human cases. We estimated that *β*_*WH*_ = *R*_0_/(*N* * *D*), where *N =* 4002, *D =* 3 and *R*_*0*_ = 0.83, based on an influenza (pH1N1) outbreak on a swine research farm in Alberta, Canada (56). We assumed that the four infected workers were responsible for the first 10 swine influenza cases. In baseline simulations, we assumed that a single infected pig was introduced when the last room was filled on Day 42.

Lastly, we assumed there was no mortality in the infected pig population. This assumption was chosen to simulate a worse-case scenario in which each hog would remain alive throughout the entire infectious period in order to better assess the impacts of vaccination and non-pharmaceutical interventions.

### Interventions

We tested four categories of interventions: mass swine vaccination, quarantine of infectious pigs, a directional workforce routine, and improvement in worker PPE. For vaccination, we assumed that vaccine efficacy (%) corresponded to the percentage of the pig population that was completely protected from both infection and ongoing transmission (i.e. they remained in the recovered compartment) (63).

In order to simulate quarantine, we incorporated four quarantine compartments (one for each room). We assumed workers adhered strictly to PPE and N95 respirator use when working with these quarantined animals and that these animals were visited only after all other pig rooms each day; therefore, we assumed no pig-to-human transmission and no between-room transmission from quarantined pigs to other pigs.

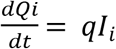, where *q* is the rate at which infectious swine move from each room to the quarantine pen.

In order to simulate the directional workforce routine, we assumed that each worker always moved from the youngest pig cohort (the pigs that entered the farm most recently) to the oldest cohort of pigs. Under this assumption, the incidence attributed to between-room transmission is shown in **Table 2** (The baseline equations were modified accordingly.)

**Table 2.** Incidence attributed to between-room transmission assuming workers work in a single direction from youngest (room 4) to oldest (room 1) pigs.

| Room Numbers | Equations |
| --- | --- |
| 4 | $\beta_N S_4(0)$ |
| 3 | $\beta_N S_3(I_4)$ |
| 2 | $\beta_N S_2(I_3 + I_4)$ |
| 1 | $\beta_N S_1(I_2 + I_3 + I_4)$ |

We also evaluated the impacts of improved PPE on reducing transmission from the workforce to pigs. We assumed workers wore either no facial covering, surgical masks, or N95 respirators and assumed perfect adherence. We assumed that the % effectiveness of masks corresponded to a % reduction in the effective contact rates from workers to swine and/or from swine to workers (64). Surgical masks are designed to trap secretions from the wearer and therefore reduce transmission from the individual, and were assumed to reduce only *β*_*WH*_ (65). N95 masks were assumed to reduce *β*_*WH*_ and *β*_*QHW*_(65).

### Simulations

All analyses and models were run in R version 4.1.0 (66) using the GillespieSSA (53) package and tidyverse (67) for plotting. All control measures were compared to the baseline model to determine their impact on total number of infected pigs per production cycle, probability of at least one workforce infection per production cycle, and time from infected pig introduction to first workforce infection. Lastly, we compared different combinations of control measures. The baseline and intervention models were run for 5,000 and 10,000 iterations to confirm that the key results were stable.

### Sensitivity analyses

To test the effects of aleatory uncertainty arising from the stochastic process implemented in our model, the coefficient of variation was calculated for total infected pigs and days to first workforce infection. To explore the impacts of variability and epistemic uncertainty, we performed a sensitivity analysis. For our sensitivity analyses, we implemented a deterministic approach using Monte Carlo sampling (5000 samples) of distributions of values for each of the model parameters (**Table 1**). We compared results from the sensitivity analysis (deterministic models) to results of our stochastic models for single control measures only (**Supplemental Table 1**).

## Results

Our baseline model (no vaccination, no quarantine, no workforce routine changes) showed that in >50% of iterations, almost the entire pig population was infected after the introduction of single infected pig within the last room to be populated, on day 42 of the production cycle (**Table 3**). The number of infected pigs peaked on average on day 61(95% uncertainty interval 42 – 65.1) of the production cycle (**Figure 2**). The probability of at least one workforce infection per production cycle was 0.61 with the first workforce infection occurring on average within 21 days of infected pig introduction (median 20.7 [range 11.8 – 33.1]; 5,000 iterations).

**Table 3.** Simulated influenza transmission dynamics under separate control measures.

| **Control Measure (CM)** | **Scenarios** | **Total Pigs Infected: Median [95% Uncertainty interval] (CV)** | **Probability of Workforce infection** | **Days from first infected pig to first Workforce Infection: Median [95% Uncertainty interval] (CV)** |
| --- | --- | --- | --- | --- |
| Baseline Model | No control measures | 3958 [1 – 3972] (0.33) | 0.61 | 20.7 [11.8 - 33.1] (0.31) |
| Mass vaccination of pigs prior to arrival on farm. | VE = 40% | 2363 [1 - 2375] (0.45) | 0.42 | 28.4 [15.1 - 44.0] (0.31) |
|  | VE = 80% | 23.5 [1 - 635] (1.09) | 0.07 | 64.2 [24.9 - 119] (0.44) |
| Quarantine of infectious pigs up to a maximum of 10 pigs | 1/q = 2 days | 3946 [1 - 3966] (0.73) | 0.43 | 25.4 [14.9 - 38.6] (0.28) |
|  | 1/q = 1 day | 3 [1 - 3961] (1.22) | 0.27 | 29.5 [17.4 - 47.3] (0.23) |
| Directional Workforce Flow from Room 4 to Room 1 only. | Intro: Room 4 | 3957 [1 - 3972] (0.33) | 0.62 | 20.8 [11.9 - 33.0] (0.31) |
|  | Intro: Room 3 | 2976 [1 - 3968] (0.35) | 0.51 | 20.9 [11.6 - 33.4] (0.32) |
|  | Intro: Room 2 | 1988 [1 - 2982] (0.39) | 0.37 | 19.7 [10.8 - 31.9] (0.32) |
|  | Intro: Room 1 | 997 [1 - 1984] (0.45) | 0.22 | 17.3 [10.1 -28.0] (0.32) |

Room number denotes which room the infected pig was introduced into, with room four being the room with the youngest pigs (filled last) and room one being the room with the oldest pigs (filled first). CV = Coefficient of Variation; VE = Vaccine efficacy; 1/q = mean time until an infectious pig is quarantined; Intro = room to which infected pig is introduced.
Days from first infected was calculated using only iterations in which a workforce infection occurred.

**Figure 2.**
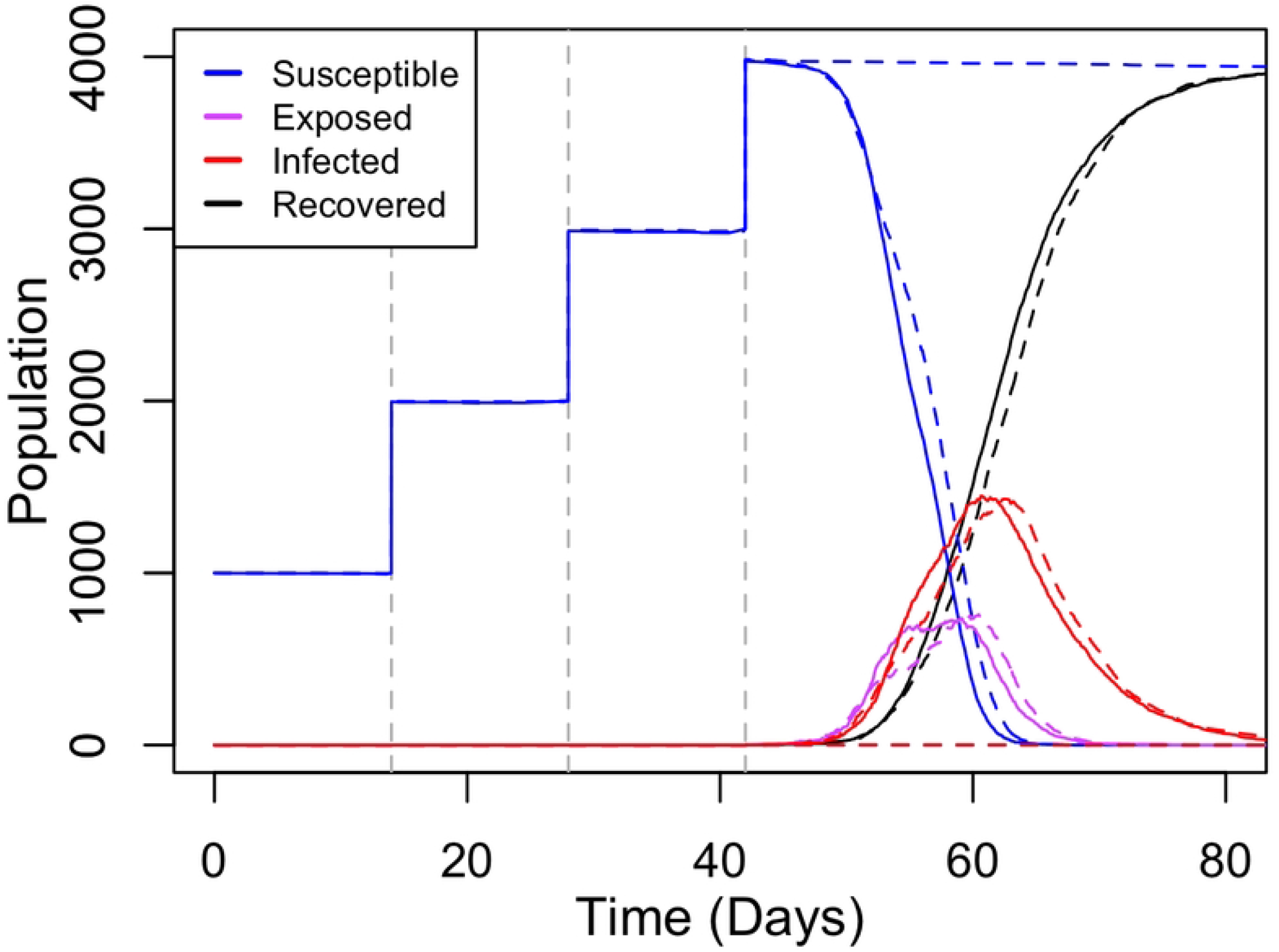
Influenza dynamics upon introduction of single infected hog on day 42. Baseline scenario - unvaccinated pigs, assuming density dependent transmission. 4,000 total hogs on the farm in 4 rooms populated 2 weeks apart. Solid lines show the median days to peak infection was 61 days (42 – 65.1) amongst 5000 iterations. Dashed lines represent the 95% uncertainty interval.

Assuming that the swine vaccine that was 40% efficacious, we found that the median total number of infected pigs was reduced from 3958 [1-3972] to 2363 [1-2375] (**Table 3**). Assuming 80% efficacy, the median total number of infected hogs was further reduced to 23.5 [1-635] (**Table 3**). The probability of workforce infection was reduced from 0.61 in the baseline model to 0.41 and 0.07 for a 40% or 80% effective influenza vaccine, respectively (**Figures 3A, 3B**).

**Figure 3.**
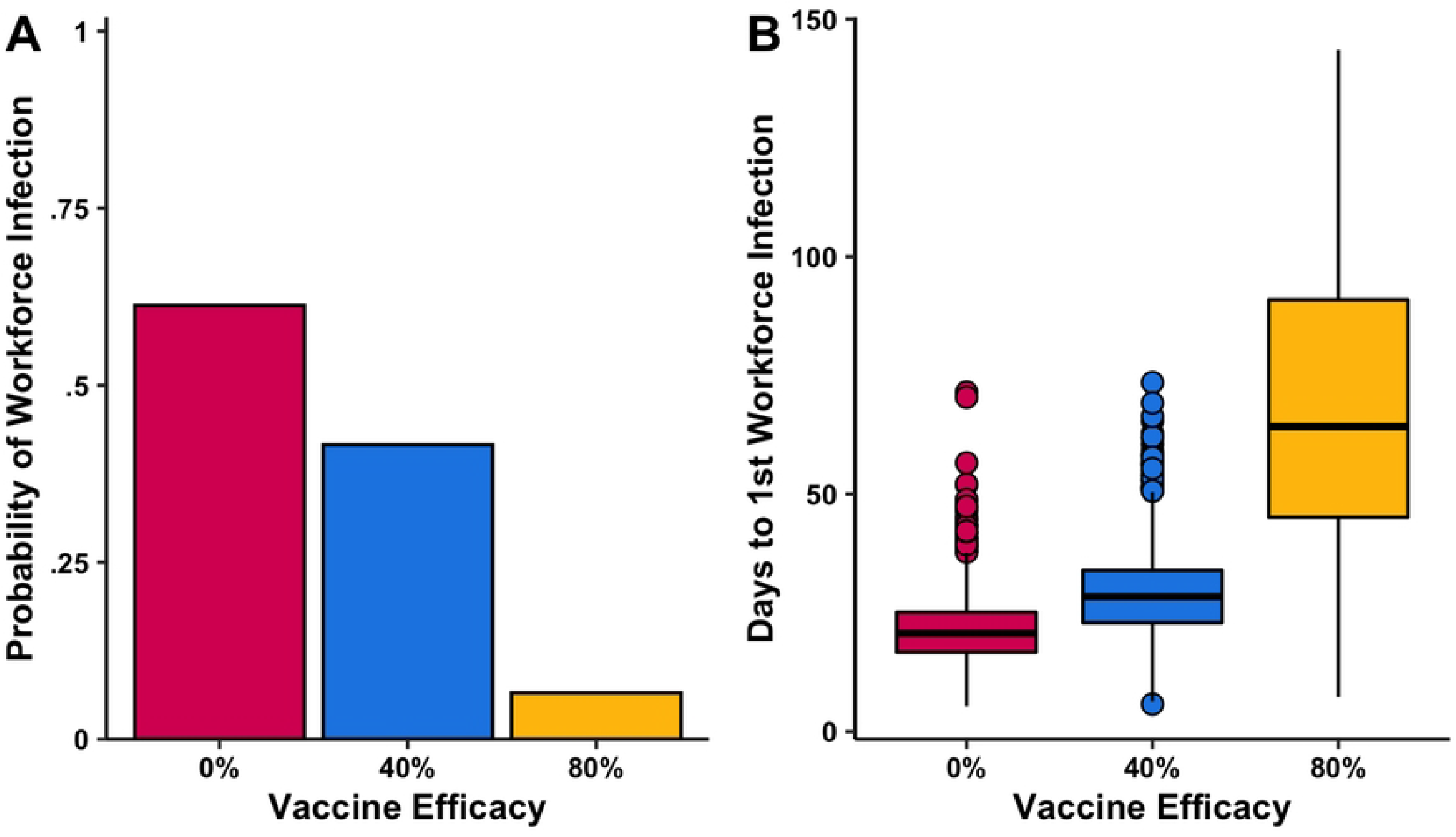
A)Probability of at least one workforce infection during one production cycle with mass vaccination of hog population and B) days from first infected pig introduction to first workforce infection with mass vaccination of hog population. Assumed introduction of a single infectious pig on day 42. density dependent transmission. 4,000 total hogs on the farm in 4 separate rooms.

When infected pigs were quarantined 1 day after they were infectious (q= 1) the model predicted a large reduction in the total number of infected pigs (3 [1 – 3966]) compared to the baseline model (3958 [1 – 3972]). There was a minor decrease in the total number of infected pigs (3946 [1 – 3966]) compared to the baseline model (3958 [1 – 3972]) when quarantining pigs at a more realistic two days after becoming infectious (q= 0.5) (**Table 3**). Quarantining infectious animals also had a modest impact on the probability of workforce infection and time to first workforce infection compared to baseline (**Table 3** and **Figures 4A, 4B**).

**Figure 4.**
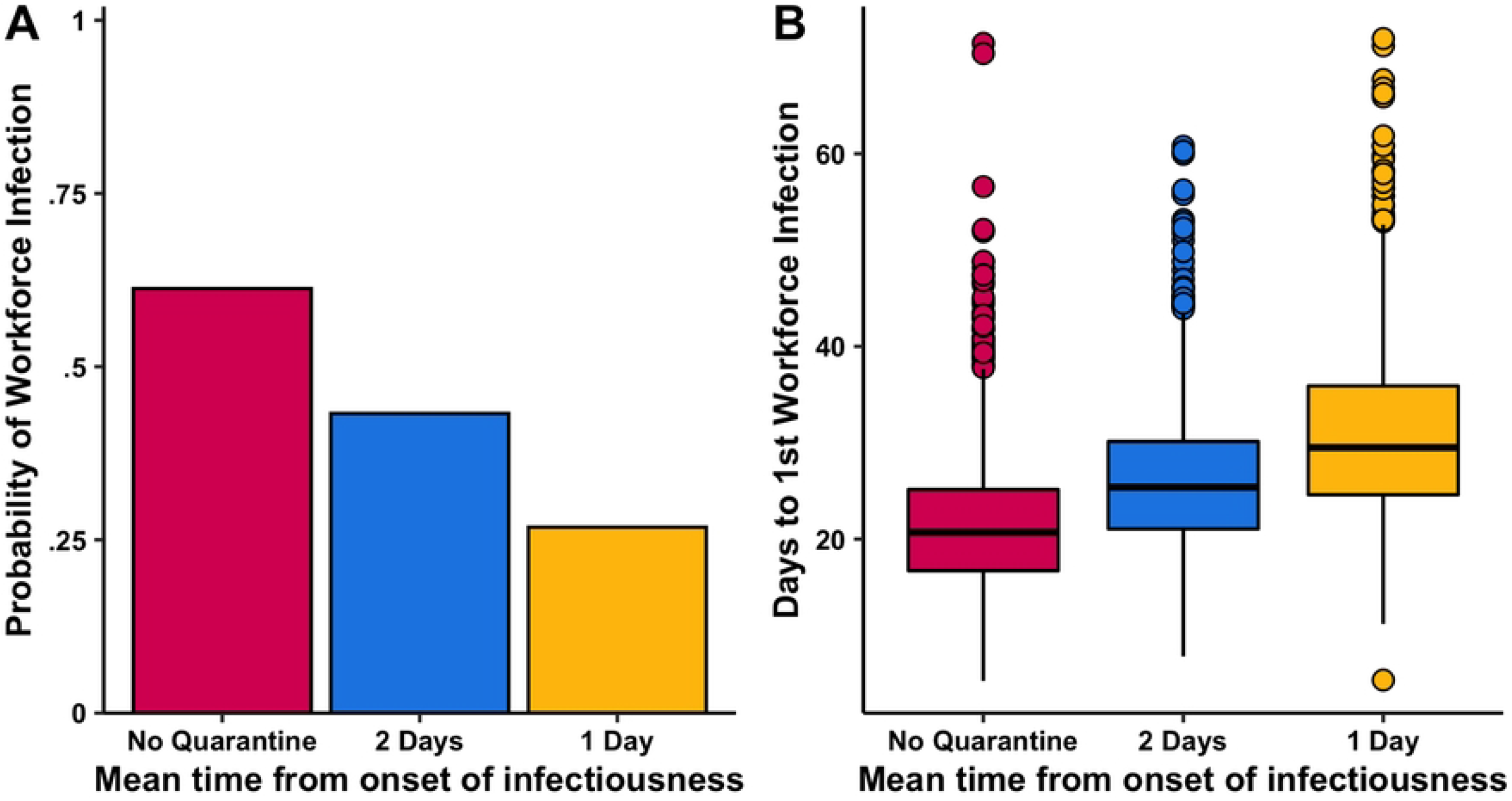
A) Probability of at least one workforce infection per production cycle with quarantine of infectious hogs and B) days from first infected pig introduction to first workforce infection with quarantine of infectious hogs. Assumed introduction of a single infectious pig on day 42, density dependent transmission and a maximum quarantine population of 10 pigs per room. 4,000 total hogs on the farm in 4 separate rooms.

Assuming workers move in a single direction from youngest pig batches to oldest pig batches led to a decreased median number of infected pigs (**Table 3**). When the infected pig was introduced in the youngest batch (Room 4), we found no significant change from the baseline scenario when comparing total infected pigs, probability of workforce infection, and time to first workforce infection (**Table 3**). However, if the infected pig was introduced earlier into the simulation in one of the older batches (Room 1 or Room 2 for example), we found a reduction in the median total number of infected pigs (Room 1 introduction: 997 [1 – 1984]; Room 2 introduction: 1988 [1 – 2982]) compared to baseline (3957 [1 – 3972]). Our models also indicated that the probability of workforce infection (**Figure 5A**) was reduced and time to first workforce infection was increased when introducing a directional workforce routine (**Figure 5B**).

**Figure 5.**
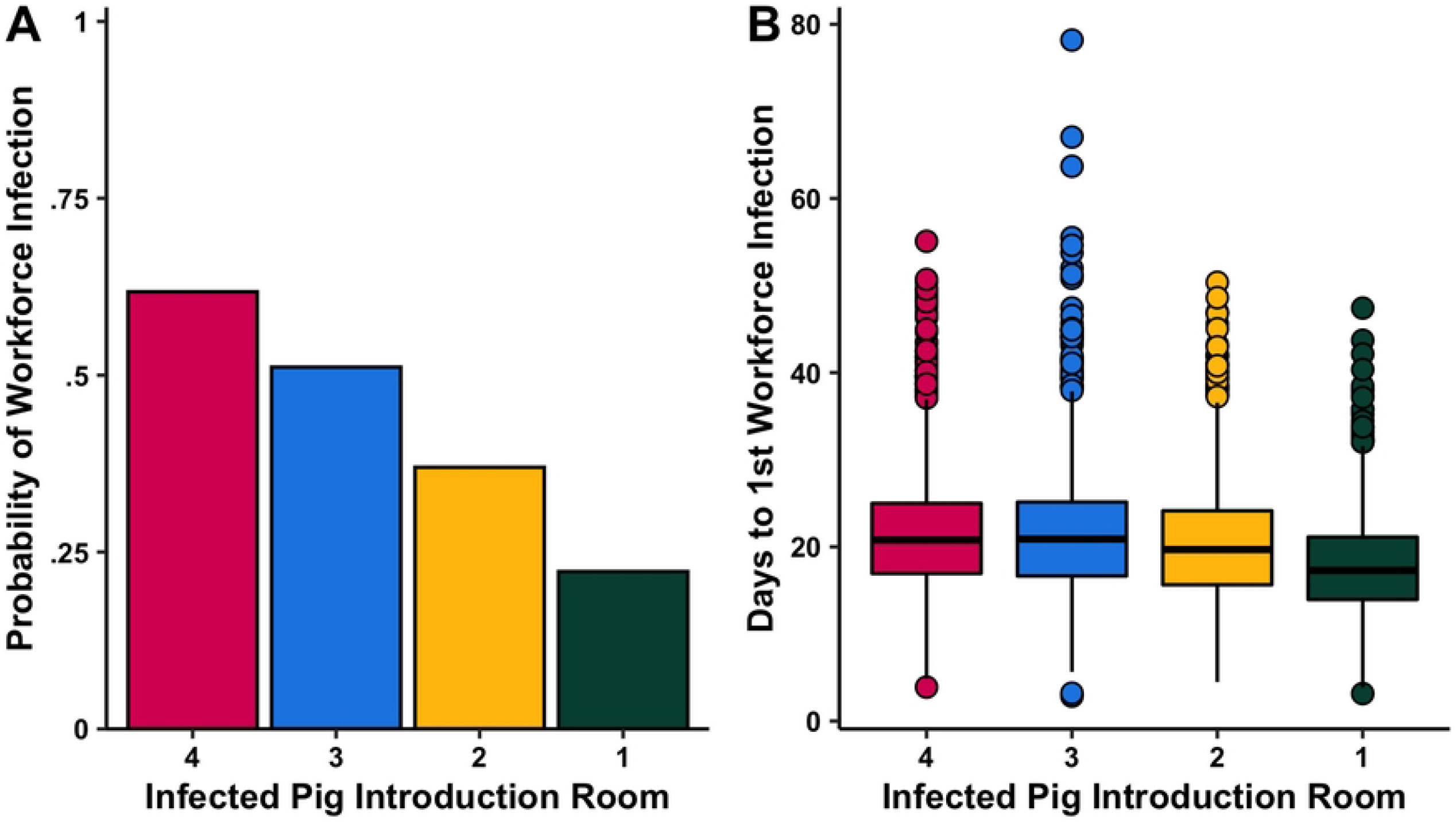
A) Probability of at least one workforce infection per production cycle with a directional workforce routine and B) days from first infected pig introduction to first workforce infection with a directional workforce routine. Assumed introduction of a single infectious pig on day 42, density dependent transmission and strict adherence of workforce working from younger pig batches to older batches. 4,000 total hogs on the farm in 4 separate rooms.

### Results of Combination of control measures

When combining quarantine of infected pigs with mass vaccination (with 40% Vaccine Efficacy), the model predicted a reduction from baseline to 2015 [0 – 2124] total infected pigs (**Table 4**). Furthermore, the probability of a workforce infection was reduced from baseline to 0.21 per production cycle in which a single infected pig was introduced. There was also an increase in time from first infected pig to first workforce infection to 48.9 days [26.1 – 77.5].

**Table 4.**
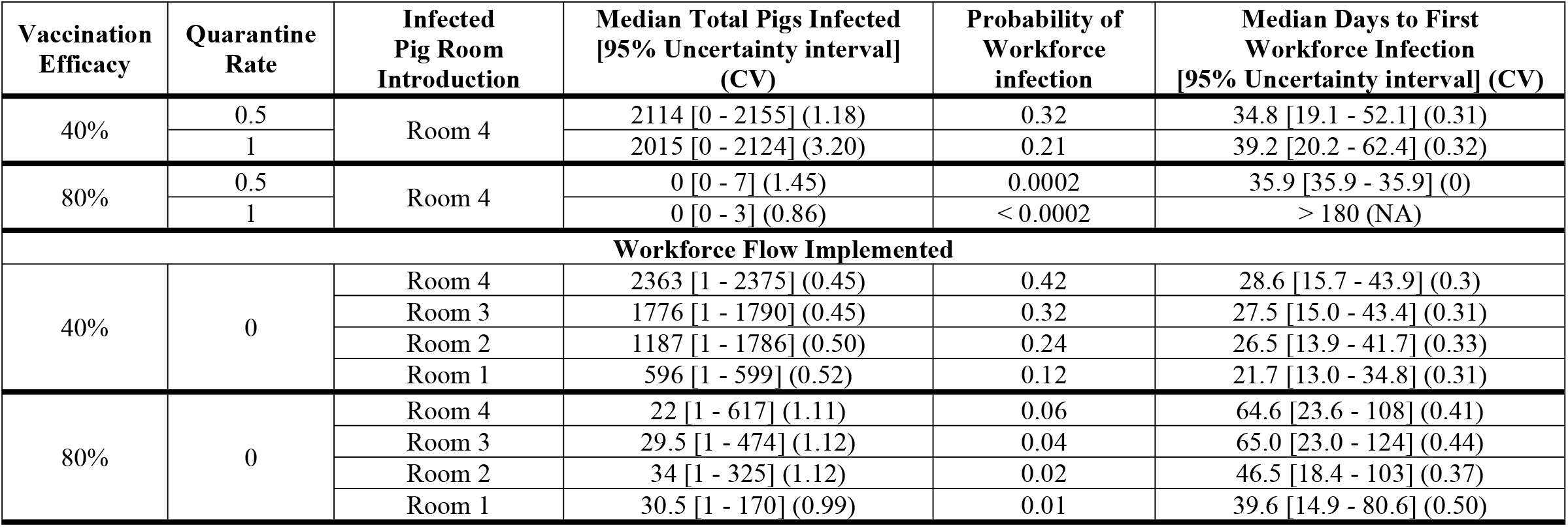
Simulated influenza transmission dynamics under combined control measures.

The model predicted additional benefits when combining a directional work routine with mass vaccination using a 40% efficacious vaccine (**Table 4**). In the best-case scenario, when the infected pig is introduced with the oldest pigs, only 596 [1 – 599] total pigs were infected. This combination of interventions also reduced the probability of workforce infection to 0.12 and increased the time to first workforce infection to 21.7 [13.0 – 34.8] compared to the baseline model (probability of workforce infection = 0.61; time to first workforce infection = 20.7 days [11.8 – 33.1]).

When all control measures were implemented (quarantine of infected pigs, mass vaccination, and workforce operation) we found that the probability of workforce infection decreased to below 0.05 per production cycle, even when the infected pig was introduced in the youngest batch of pigs (Room 4) and with a 40% vaccine efficacy (**Table 5**). The probability of workforce infection was reduced to <0.0002 when vaccine efficacy was improved to 80%.

**Table 5.** Simulated influenza transmission dynamics when all control measures are implemented.

| **Vaccination Efficacy** | **Quarantine Rate** | **Infected Pig Room Introduction** | **Total Pigs Infected [95% Uncertainty interval] (CV)** | **Probability of Workforce infection** | **Days to first Workforce Infection [95% Uncertainty interval] (CV)** |
| --- | --- | --- | --- | --- | --- |
| 40% | 1 | Room 4 | 1 [1 - 2333] (3.24) | 0.03 | 51.2 [26.2 - 88.6] (0.36) |
|  |  | Room 3 | 1 [1 - 1755] (3.10) | 0.03 | 44.8 [26.0 - 7.4] (0.32) |
|  |  | Room 2 | 2 [1 - 1174] (3.17) | 0.02 | 45.2 [22.5 - 71.9] (0.36) |
|  |  | Room 1 | 2 [1 - 591] (3.00) | 0.01 | 31.6 [22.1 - 50.3] (0.30) |
|  | 0.5 | Room 4 | 4 [1 - 2359] (1.19) | 0.18 | 40.3 [22.1 - 62.9] (0.31) |
|  |  | Room 3 | 3 [1 - 1774] (1.23) | 0.14 | 39.4 [20.6 - 61.5] (0.34) |
|  |  | Room 2 | 4 [1 - 1186] (1.21) | 0.09 | 36.2 [19.5 - 62.8] (0.37) |
|  |  | Room 1 | 4 [1 - 596] (1.24) | 0.06 | 30.3 [19.4 - 48.8] (0.31) |
| 80% | 1 | Room 4 | 1 [1 - 4] (0.86) | <0.0002 | > 180 days (NA) |
|  |  | Room 3 | 1 [1 - 4] (0.86) | <0.0002 | > 180 days (NA) |
|  |  | Room 2 | 1 [1 - 4] (0.81) | <0.0002 | > 180 days (NA) |
|  |  | Room 1 | 1 [1 - 4] (0.87) | <0.0002 | > 180 days (NA) |
|  | 0.5 | Room 4 | 1 [1 - 8] (1.47) | 0.0002 | 36.7 [36.7 - 36.7] (0) |
|  |  | Room 3 | 1 [1 - 8] (1.48) | <0.0002 | > 180 days (NA) |
|  |  | Room 2 | 1 [1 - 8] (1.49) | <0.0002 | > 180 days (NA) |
|  |  | Room 1 | 1 [1 - 8] (1.45) | 0.0002 | 9.29 [9.29 - 9.29] (0) |

### Improved worker PPE

To test the effects of worker PPE and adherence to PEE mandates on influenza transmission from pigs to workers, simulations were performed using the same assumptions and parameters as the baseline model. We found that when a worker wore a surgical mask the probability of pig-to-worker transmission was unchanged from 0.61. However, when workers adhered to strictly wearing an N95 mask, the probability of pig-to-worker transmission was reduced to 0.08. We also performed simulations to determine the impacts of a worker infected with influenza virus (swine associated H1N1, H1N2 or H3N2) infecting the pig population on our growing unit. The simulations found that the probability of at least one pig being infected from an infected worker was 0.54. Our results also showed that this probability was reduced to 0.28 when workers wore a surgical mask and 0.10 when wearing an N95 mask, assuming the effective contact rate was reduced by the same percentage as published effectiveness values (64), and workers had perfect adherence to facial covering policies. We also found that the median number of pigs infected during these outbreaks was 3936 [0 – 3967] for no mask, 0 [0 – 3977] when workers wore a surgical mask, and 0 [0 – 3968] when workers wore N95 masks.

## Discussion

Our research aimed to simulate interspecies influenza transmission on a typical US indoor pig growing unit in order to test the effects of vaccination and non-pharmaceutical interventions, alone and in combination. To evaluate the effects of these interventions, we developed a baseline model where no vaccination was implemented, no pigs were quarantined, and workers mixed randomly with the pigs. Upon introducing a single infected pig with a single subtype of influenza virus, the model predicted an outbreak in 90% of our iterations that infected almost the entire pig population (more than 3000 pigs infected) on the farm. Our findings were consistent with empirical data supporting widespread virus transmission in swine farms, and consistent with modeling studies on different types of swine farms (46, 47, 49, 68). However, the effective contact rate parameters in the model were based on relatively few empirical studies of different subtypes (all associated with swine). We did not simulate the transmission of subtypes typically associated with humans or avian species.

Assuming mass vaccination of swine with 40% efficacy, which is typical of vaccines currently in use, the median number of infected pigs was reduced by 1595 to 2362 [0 - 2374]. Furthermore, the probability of workforce infection was reduced by 0.2 to 0.4. Reducing the likelihood of transmission to the workforce can help reduce the likelihood of influenza of swine origin spreading to the local community, as well as ensuring a healthy workforce (69).

Overall, our results suggest that vaccination is beneficial in the control of endemic influenza, even with suboptimal efficacy. However, in cases of a novel influenza subtype, a vaccine may not be readily available. In these cases, non-pharmaceutical interventions were also predicted to be a modestly effective strategy to mitigate transmission. Our results indicated that quarantining symptomatic animals has some efficacy when applied in the absence of other control measures. However, its impact was assumed to be limited by the number of quarantine pens available. Our predictions were also based on several other important assumptions: we assumed that quarantine pens and biosecurity measures practiced were sufficient to prevent any onward transmission from quarantined pigs to other pigs or humans, and that pigs were quarantined on average 1 or 2 days after the onset of infectiousness.

In some circumstances it may be more practical to implement a revised workforce routine, in which workers always work from the youngest pig batches first and move sequentially to the batches with older pigs. The model predicted up to 50% reduction in the number of infected pigs compared to the baseline model, depending on which the room the infected pig is introduced to. Further, adherence to the directional work routine greatly reduced the likelihood of the workforce being infected. This directional work routine could be further improved by workers entering rooms with known infected pigs last of all, each day. Our results also indicated that strict adherence to wearing an N95 mask greatly reduced the probability of workers becoming infected from sick pigs. This is not surprising as N95s are designed to protect the wearer from infection, and if fitted properly, they can reduce transmission as well, while surgical masks are designed to trap secretions from the wearer and therefore reduce transmission from that individual (65). These results suggest in the face of a novel influenza subtype, strict adherence to new worker routines and wearing N95s will likely reduce the likelihood of pig-to-worker and worker-to-pig transmission.

We assumed that there were only two workers on a farm this size, which is typical of intensive indoor hog production in the US. It is likely that human infections would be more common on farms with more workers, in which case modelling efforts should include worker-to-worker transmission. Further, influenza in swine peaks in spring and fall months, and therefore, probability of worker infection varies according to season (17).

We found that for most model simulations, the CV for total infected pigs and days to first workforce infection was less than one. However, our findings also suggest that more variability was introduced (CV > 1) when we implemented multiple control measures in combination.Increasing the number of simulation iterations would likely lead to reduced CV values. Our sensitivity analysis (deterministic approach with Monte Carlo sampling for transmission parameters, latent and infectious periods) produced similar results (**Supplemental Table 1**) to our stochastic model with the exception of workforce infections (**Table 3**).

We focused on transmission of a single subtype of influenza within a single production cycle on a single US indoor growing unit. We did not evaluate the likelihood of between production cycle infections that could arise from improper disinfection within swine housing facilities, equipment, or transportation vehicles. Results from this study may not apply to outdoor units, facilities that are not grower specific, and swine practices outside of the US. We assumed no prior acquired immunity due to infection before pigs entered the growing facility. There were important knowledge gaps in the reviewed literature, which led to uncertainty in effective contact rate values. We based our transmission parameters from single experiments with a mixture of subtypes. Importantly, due to limited availability of empirical data the between-room transmission parameter combines both fomite and aerosol transmission. Interventions targeted at reducing aerosol transmission or fomite transmission may have very different impacts on between-room transmission, and these differences may also be sensitive to the farm building structure. However, despite these uncertainties our findings demonstrate the potential importance of influenza transmission between workers and pigs on US swine farms and demonstrate that relatively practical interventions could be applied to reduce transmission in order to mitigate the economic and public health impacts, especially in the face of a novel subtype. Further work is needed to test the acceptability of these measures to workers and swine farms, and to further understand the role that this swine-human interface has on the dynamics of influenza generally.

---

**Supplemental Table 1 Sensitivity Analysis Results**

*to calculate days to first Workforce infection and the deterministic nature of these models we only used iterations in which to total infected workforce count exceeded 0.5

## Notes

### Competing Interest Statement

The authors have declared no competing interest.

